# The iron maiden. Cytosolic aconitase/IRP1 conformational transition in the regulation of ferritin translation and iron hemostasis

**DOI:** 10.1101/2021.07.28.454104

**Authors:** Cécilia Hognon, Emmanuelle Bignon, Guillaume Harle, Nadège Touche, Stéphanie Grandemange, Antonio Monari

## Abstract

Maintaining iron homeostasis is fundamental for almost all living being, and its deregulation correlates with severe and debilitating pathologies. The process is made more complicated by the omnipresence of iron and by its role as a fundamental component of a number of crucial metallo proteins. The response to modifications in the amount of the free iron pool is performed via the inhibition of ferritin translation by sequestering consensus messenger RNA (mRNA) sequences. In turn this is regulated by the iron-sensitive conformational equilibrium between aconitase and IRP, mediated by the presence of an iron-sulfur cluster. In this contribution we analyze by full-atom molecular dynamics simulation, the factors leading to both the interaction with mRNA, and the conformational transition. Furthermore, the role of the iron-sulfur cluster in driving the conformational transition is assessed by obtaining the related free energy profile via enhanced sampling molecular dynamics simulations.

## 1. Introduction

Iron is a fundamental element, which is involved in key biological processes [1–4], such as oxygen transport in hemoglobin, electron transfer, muscle contraction, and serves as a co-factor for many enzymes. As a matter of fact, a number of iron-containing protein has been identified, and the latter are involved in extremely diverse functions [5]. For instance, Fe-S clusters have been assigned both a structural and redox role in DNA repair proteins and other enzymes [6–8]. In addition to protein-embedded iron, a limited quantity of free iron is also available in cells, in what is called free iron pool (FIP) [9–11]. FIP is constituted by iron in both II and III oxidation states and the metal is usually complexed with different, and not yet very well characterized, ligands and chelatants [12]. Despite its limited characterization, FIP is essential for cells viability, and its dysregulation may be correlated with cellular instability, and at a systemic level, with the development of debilitating pathologies, including cancer [13–16] and mitochondria failure [17,18]. On the other hand, FIP iron accumulation, and especially the accumulation of Fe^2+^ is also inducing considerable oxidative stress, most notably due to the production of peroxyl radical via Fenton reaction [19]. As a logic consequence cells have developed complex regulatory machineries to stabilize and regulate the amount of FIP [20,21].

It is recognized that the regulation of iron homeostasis involves tuning the expression of several genes involved in storage, outtake and intake of iron such as ferritin H and L (involved in iron storage), ferroportin (involved in iron outtake) and transferrin receptor (TFR1 involved in iron uptake) [22,23]. These regulations have to be monitored rapidly to react to all changes in iron availability. Thus, cells possess a post-transcriptional regulatory mechanism that involves the binding of messenger RNA (mRNA) consensus sites present in untranslated sequences (UTR) of the iron regulatory genes by specific proteins [24–28].

Indeed, in presence of high iron level, cytosolic aconitase (cAco) embeds a Fe-S cluster which confers an enzymatic activity catalyzing the isomerization of citrate in cis-citrate via cis-aconitate. Conversely, when the FIP level decreases cAco loses the Fe-S cluster triggering a conformational transition that is accompanied with the loss of the catalytic activity [29,30]. Furthermore, in its iron-free conformation the protein, known as iron regulating protein 1 (IRP1), is able to bind a hairpin mRNA sequence, the iron regulating element (IRE) which is highly conserved and present in the ferritin, ferroportin and transferrin receptor transcripts of vertebrates, while it is absent in non-vertebrate despite the conservation of the cAco/IRP1 equilibrium [31–33]. The binding of IRP1 on IRE, located in 5’UTR, obviously induces a blockage of ferritin and ferroportin translation, leading to their downexpression and ultimately to the increase of FIP level (Scheme 1) [34,35]. On the contrary, the binding of IRP1 on the IRE present in the 3’UTR of the transferrin receptor leads to mRNA stabilization and thus to an elevation of its translation allowing an increase in iron uptake. Once the cellular iron concentration has risen again sufficiently, the equilibrium between cAco and IRP1 is altered releasing IRE and reinstating the translation of these genes.

**Scheme 1.**
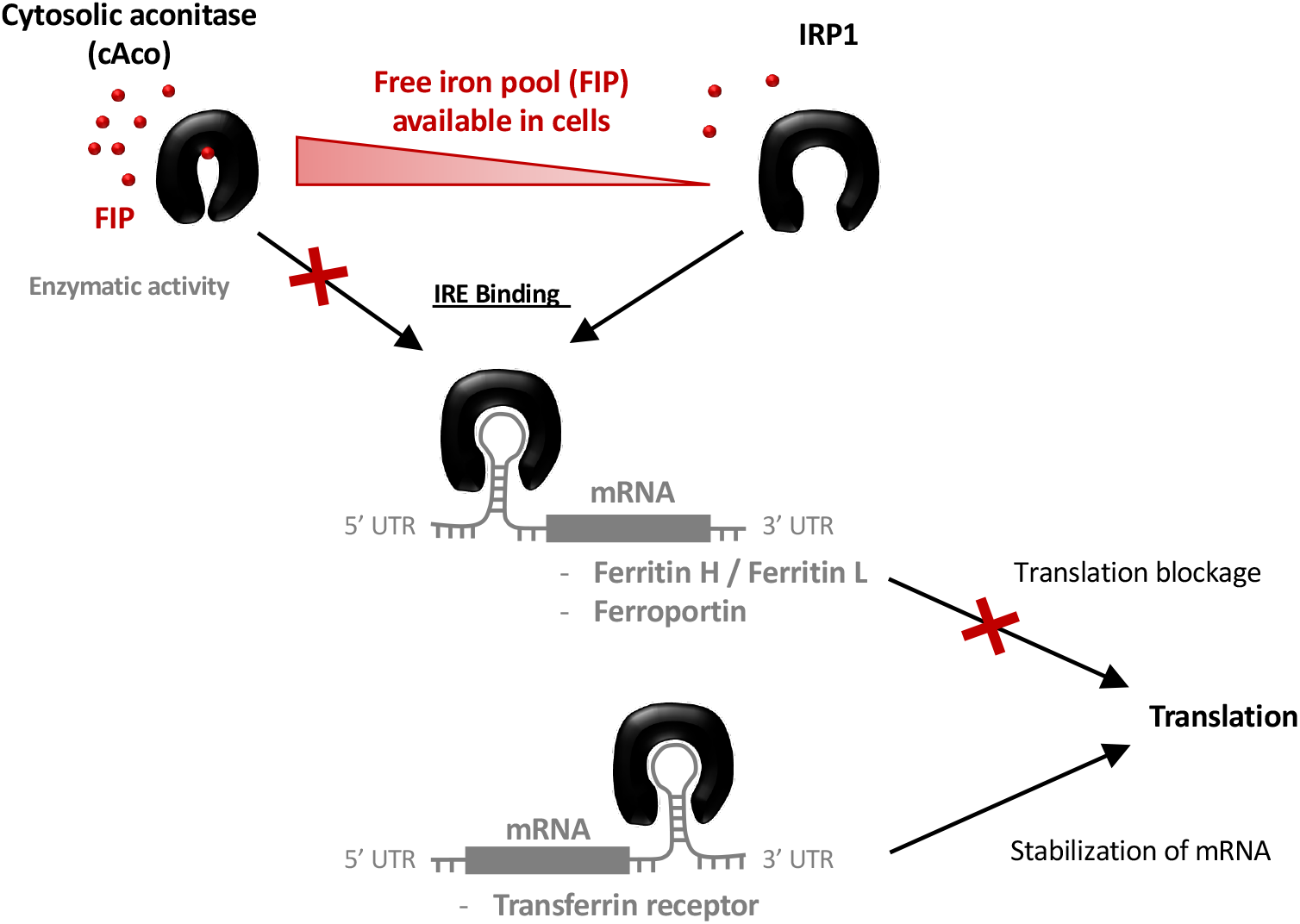
Mechanism of the regulation of FIP level by the cAco/IRP1 conformational equilibrium and the binding with IRE mRNA.

From a structural point of view, different crystal structures of cAco and IRP1, also complexed with the IRE hairpin sequence, have been resolved [7,36]. Evidencing that the conformational translation responsible for homeostasis regulation can be summarized as the passage from a closed (cAco) to an open (IRP1) structure which exposes the RNA binding motifs. However, some important questions still hold concerning the molecular bases driving the mechanisms. In particular the role of the Fe-S cluster in stabilizing the cAco form has not been clearly evaluated, as well as the reasons leading to the selective interaction with IRE consensus sequence. Furthermore, the flexibility and intrinsic dynamic of IRP1 bound and unbound to RNA is also lacking and could provide important insights to clarify its mechanism of action.

In this contribution we resort to all atom classical molecular dynamic (MD) simulations to explore the behavior of cAco, IRP1, and IRE and their mutual interactions using the consensus IRE of the ferritin transcript as a model. Furthermore, we also provide enhanced sampling MD simulation to assess the free energy profile for the cAco IRP1 transition in presence and in absence of a Fe-S cluster, hence assessing the thermodynamics factors at the base of the first step of iron hemostasis.

The deregulation of the cellular iron level is associated to a series of debilitating pathologies, including cancer, yet its current comprehension, from a structural point of view, is still far from being ideal. Unraveling the key biophysical effects behind the complex structural transitions taking place may lead, in the future, to the development of novel therapeutic strategies, also involving therapeutic mRNA protocols.

## 2. Materials and Methods

The initial structure of the IRP1/IRE complex has been obtained from pdb code 3snp [7], the initial conditions for the MD simulations of the isolated IRP1 and IRE moieties have been obtained from the same crystal structure. The structure of the human cAco enzyme has been instead retrieved from the pdb code 2b3y [36]. Even if the two structure are referring to different organisms the sequence homology is exceeding 95% hence assuring meaningful comparison.

For all the MD simulations the amber2014 force field has been considered [37] while in the case of nucleic acid the bsc1 corrections [38,39] to enhance the description of the backbone parameters have also been taken into account. The Fe-S (4Fe-4S) cluster has been modeled using a specifically tailored set of parameters designed by Carvalho and Swart [40,41] and already used by some of us in the modeling of endonucleases [42]. All the systems have then been solvated in a cubic TIP3P [43] water box with a 9 Å buffer and K+ cations have been added to ensure electroneutrality. MD simulations have been performed using the NAMD code [44,45] a constant temperature (300 K) and pressure (1 atm) has been enforced using the Langevin thermostat [46] and barometer [47], hence yielding a NPT simulation. After 5000 minimization steps, the system has been equilibrated for 9 ns during which constraints on the protein backbone and on the nucleic acid heavy atoms have been progressively released, and finally the production runs were performed. The equilibrium MD simulation reached 200 ns for the IRP1/IRE, IRP1, and IRE. In the case of cAco including the Fe-S cluster we also performed 200 ns, while the MD simulation of Fe-S-missing cAco was propagated for 500 ns. A time step of 4 fs has been consistently used thanks to the hydrogen mass reparation (HMR) strategy [48] in combination with the Rattle and Shake procedure [49]. Particle mesh Ewald (PME) [50] has been used to treat long-range interactions with a cut-off of 9 Å.

For the enhanced sampling procedure, we chose to sample a collective variable based on the RMSD difference (ΔRMSD) between two representative states, namely the open and closed conformations. While the closed conformation can be easily obtained from the human cAco equilibrium simulation, the open conformation has been built starting from the equilibrium MD of the chicken IRP1 after manually mutating the residue to obtain the same sequence of the human protein. Note that the stability of the mutated system has been checked and validated by an equilibrium MD. The enhanced sampling MD was performed on two situations, i.e. including or excluding the Fe-S cluster. Since the latter is not covalently bound to the cAco enzyme and to avoid its release in the medium a constraint to the distance with the Ser 435 and Cys 436 residues has been imposed. The potential of mean force (PMF) has been obtained using a combination of extended adaptative biasing force (eABF) [51] and metadynamics [52] in what is known as meta-eABF [53,54].

Most notably this choice allowed to threat the system of interest without its separation in windows. Enhanced sampling MD simulations have been performed using NAMD and its Colvar interface.

All the simulations have been visualized and analyzed using VMD [55].

## 3. Results

### 3.1. Structure of IRE mRNA

As stated in the introduction the post transcriptional regulation of ferritin expression is regulated by a well conserved mRNA motif, IRE, which assumes a hairpin sequence [56,57]. Notably, while some sequence variations in IRE exist, both between species and in the same organism, the hairpin motif appears as fundamental to assure recognition by IRP1, with a particular role played by the loop region, which is particularly well conserved [35]. In addition, some hanging uncoupled nucleobases are also present and may lead to supplementary interactions with the regulatory protein. Representative snapshots extracted from the MD simulation of a solvated IRE strand are reported in Figure 1 together with the time series of the root mean square deviation (RMSD). Two contrasting evidences may be underlined from its inspection: while globally the strand appears quite flexible, leading to a relatively high value of the RMSD (∼ 11 Å), the loop region is well conserved, as well as the global hairpin arrangement. Interestingly, the most important movement observed can be described as a global bending of the RNA strand. This in turn leads to a coexistence of straight and bent sequences, in which the loop area is approaching the double strand arrangement, and the terminal bases. Notably, two overhanging, solvent exposed, bases are also observed at the edges of the loop and in the double-strand region. If these extrahelical bases experience, as expected, an enhanced flexibility they do not induce any particular structural deformation. The global flexibility of the RNA hairpin is also confirmed by the fact that the RMSD spans a relatively large interval and experiences an oscillatory behavior. Coherently, the bending movement can also be evidenced by monitoring the distance between the hanging nucleobases U 6 (in the loop region) and U 19 (in the double helical region), which is reported in Supplementary Information and which shows a behavior strongly correlated with the RMSD inducing a sort of breathing mode.

**Figure 1.**
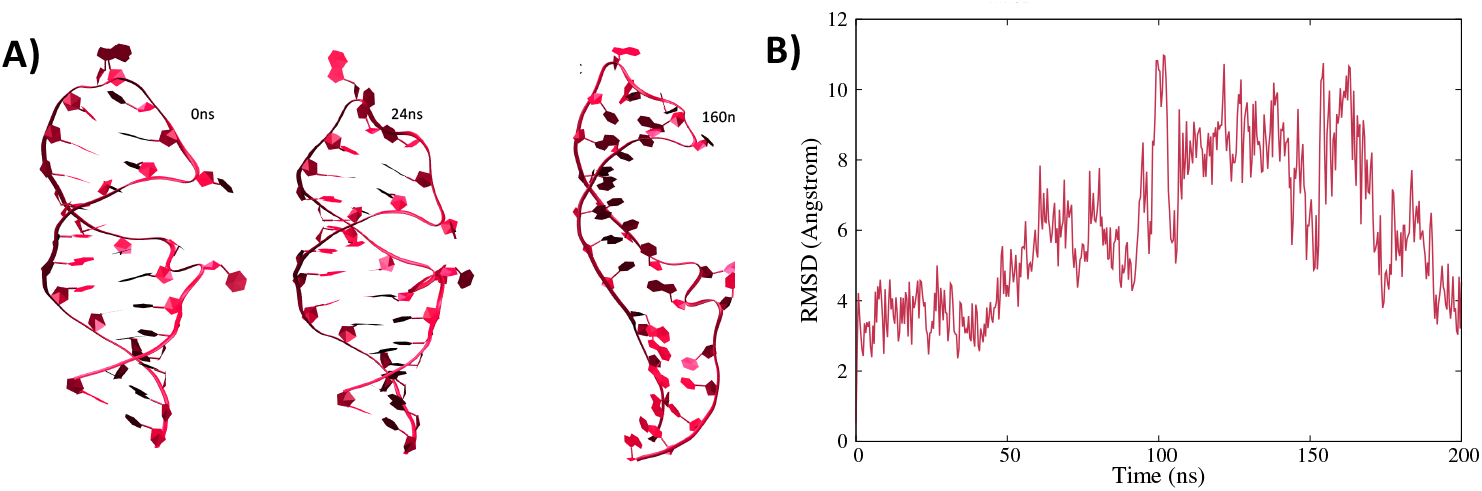
Representative snapshots extracted form the MD trajectory showing the structural evolution of the solvated IRE mRNA hairpin (A) and the corresponding RMSD (B).

### 3.2. Structure of the IRP/IRE complex

As stated in the introduction the regulation of ferritin translation is modulated by the complexation of the IRE unit performed by the IRP1 protein, i.e. the same protein sequence forming cAco but lacking the Fe-S cluster. The results of the MD simulation for the IRP1/IRE complex are reported in Figure 2, and shows that the interaction between the RNA sequence and the protein leads to the formation of a persistent and rather rigid aggregate. Interesting, while the central double-strand helical region of the nucleic acid stays rather solvent exposed, the main contact points involve the loop and the edge bases, as well as the extrahelical extruded nucleotides, mainly uracil. A striking reduction of the flexibility of the mRNA is observed upon complexation, since its RMSD now barely reaches 3 Å.

**Figure 2.**
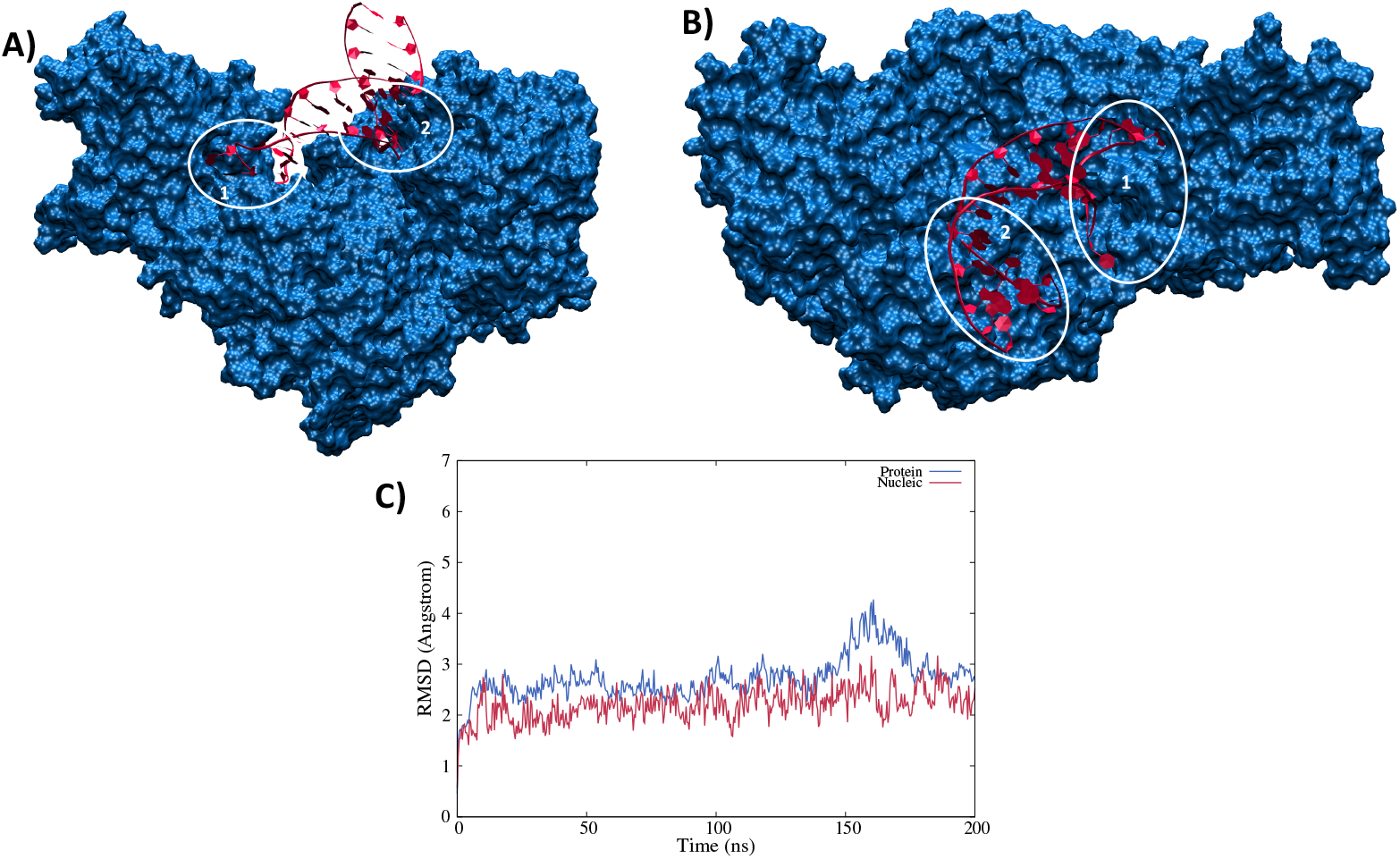
Side (A) and top (B) view of a representative snapshot of the MD simulation. The two main contact points involving the loop (1) or the edge (2) regions are highlighted in white circles and the time series of the RMSD for the protein (blue) and nucleic acid (red) are given in panel C.

The same rigidity is also observed for the protein moiety, whose RMSD is also plateauing at around 3 Å with rare peaks at 4 Å. As mentioned, two main contact points can be observed and are highlighted in Figure 4A and 4B. They involve either the loop region (1) or the edge bases of the mRNA sequence (2) and appears as extremely persistent all along the trajectory. Anchoring the RNA at the two extremities may also justify the increased rigidity of the strand. The involvement of the loop region in the direct contact with IRP1 protein is also coherent with the experimental observations pointing to the fact that this motif is, indeed, well conserved in IRE regulatory area, and is necessary to regulate ferritin translation. In both regions we may identify a complex network of amino acids interacting both with the RNA backbone and with specific and exposed nucleobases motifs as reported in Figure 3. Interestingly enough, while the prominent amino acids in the contact area are polar or basic, such as Asn, Asp, or Thr, no salt bridges involving the RNA backbone can be evidenced. This observation is in striking contrast with some of the most common protein/nucleic acid interaction patterns that involves the, rather unspecific, electrostatic binding of the negatively charged backbone of the DNA/RNA fragment. The interaction involving hydrogen bonds with specific nucleobases are also especially important in the loop region, most probably adding up to the global selectivity of the interaction.

**Figure 3.**
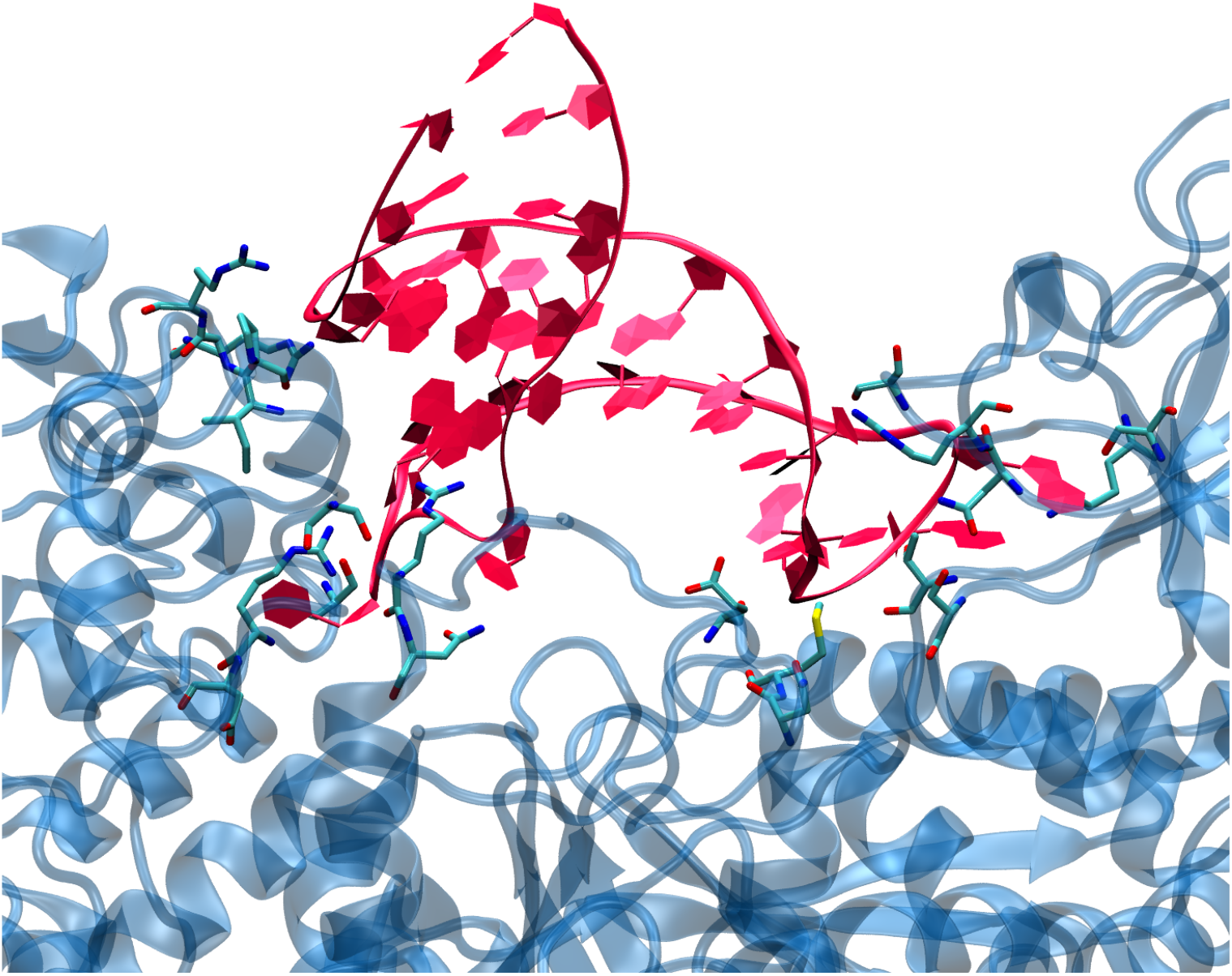
A zoom on a representative snapshot of the MD simulation of the IRP/IRE complex showing the contact region and highlighting the interacting residues.

**Figure 4.**
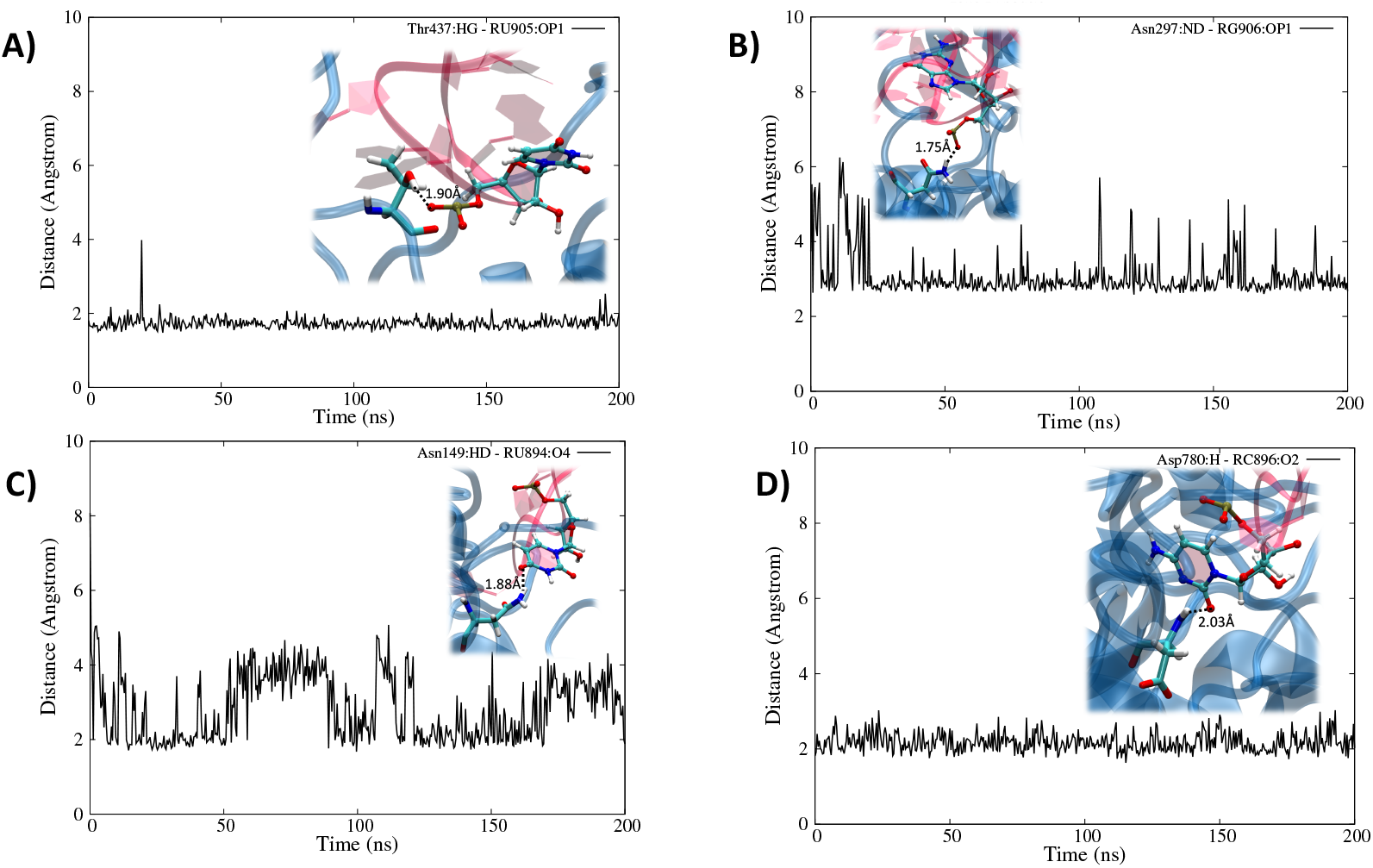
Time series of the distances between selected interacting residues involving the protein and IRE involving either the loop, Thr 437 and U (A) and Asn 297 and G (B), or the edge region, Asn 49 and U (C) and Asp 780 and C. Representative snapshots highlighting the specific interactions at atomic level are also given as inlays in each panel.

Interestingly, and as reported in SI, we may also identify some interactions which, while not present in the crystal structure, develop in the course of the MD simulation and are quite persistent, see for instance Arg 353 and Arg 148 developing salt bridges with the phosphate of guanine and uracil, respectively. This reorganization however is local and is not accompanied by any major structural deformation of either the protein or the nucleic acid; on the contrary it confirms the high stability and affinity of the IRP1/IRE complex. In Figure 4 we also report the time series of some distances characterizing chosen paradigmatic interactions involving either the loop (Figure 4A and 4B) or the edge (Figure 4C and 4D) region.

Indeed, we may also underline that the negative RNA phosphates in the backbone are mainly stabilized via the formation of hydrogen bonds, as shown in Figure 4A involving the OH group of a Thr, or the amine group of Asn in Figure 4B. Note, however, that while the interaction in Figure 4A was present already in the crystal structure and leads to extremely stable time series, the second one appears looser, showing some evidences of local and momentarily loss of the interaction. This was particularly true before equilibration, in the crystal structure the characteristic distance exceeds 6 Å and it populates values compatible with the formation of stable hydrogen-bonds only after equilibration.

Conversely, in Figure 4C and 4D we evidence interaction taking place directly with the nucleobase elements. While Figure 4C shows the interaction between the lateral chain of Asn and the electron-rich oxygen atom of uracil, Figure 4B deals with the interaction between the carbonyl group of cytosine and the peptidic NH domain of an Asp residue. This latter evidence is important since it also shows that the protein backbone is directly participating to the stabilization, which also constituting a most paradigmatic example of the prominence of Hydrogen bond patterns over salt bridges. Furthermore, while the interaction represented in Figure 4D is extremely stable, as evidenced by the time series, the more classic one in Fig 4C happens to be quite dynamic and flexible, and hence less important to assure the global stability of the complex.

As final notes one should also point out that the hanging nucleobase of IRE participate to the recognition, and that because the contact points are located at the extremity of the RNA strand, IRP1 should assume a rather extended and open conformation to accommodate and efficiently sequestrate the mRNA strand by assuming a pincer conformation.

### 3.3. Structural features of unbound IRP1 and cAco

The analysis of the equilibrium MD simulation as seen in Figure 5 confirms that the protein remains globally open and maintains the pincer-like shape. However, an increase of flexibility is evidenced upon the removal of IRE. Indeed, in the apo form IRP1 exhibits a sort of oscillatory or breathing movement characterized by the partial closing and opening of the pincer, as can be observed in Figure 5A and 5B. If globally the distance between the extremities of the pincers spans an interval of more than 25 Å, it is also evident that values below 50 Å are not observed in the course of the MD simulation. Note, as a comparison that a distance between the pincers edges of about 60 Å is observed both in the crystal structure and in the IRP1/IRE simulation. Hence, while the absence of IRE induces a global flexibilization of IRP1 significant energetic barriers prevent the protein to assume more closed conformations, which would be incompatible with mRNA recognition and sequestration.

**Figure 5.**
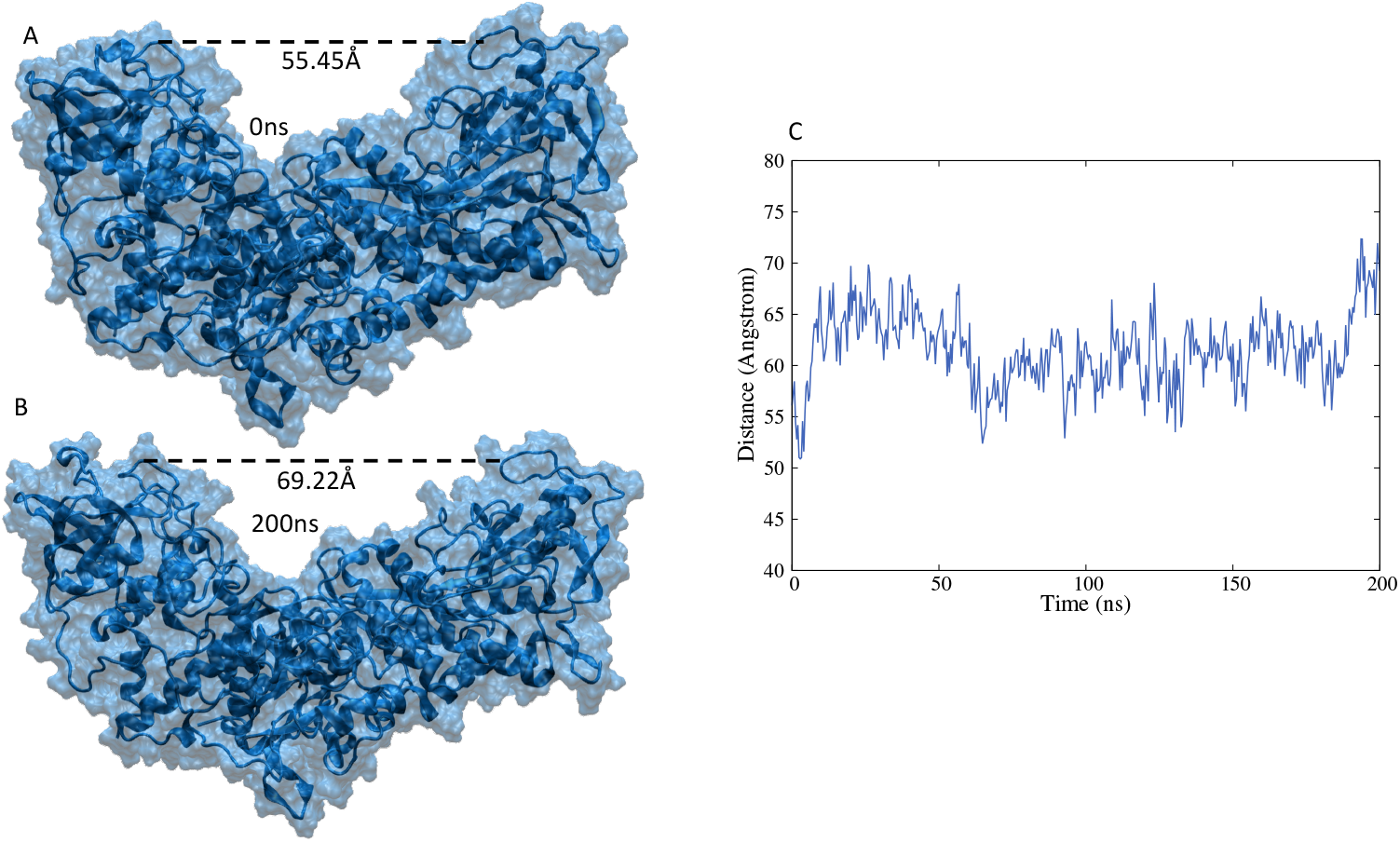
Representative snapshots of unbound IRP (A and B) and time series of the evolution of the distance between the extremities of the pincer-like domains (C).

In the case of cAco, which has the same sequence of IRP1 with the supplementary inclusion of the Fe-S cluster, the MD simulation shows a different behavior. Indeed, cAco presents a very rigid and compact structure in which the pincers are totally closed, burying the IRE recognition area in the solvent inaccessible regions (Figure 6). Note also that the Fe-S cluster is buried deep in cAco and is located at the basis of the pincers, where it interacts particularly with the residues Hys 206, Asn 297, Ser 435, Cys 436, and Asn 534, exerting a force sufficient to lock the pincers in the closed conformation. The crucial role of the Fe-S cluster can also be evidenced by the results of the MD simulation in which the former has been removed from the cAco closed conformation. In this case (Figure 6B) even though we do not recover the equilibrium open conformation of IRP1 we can note a more pronounced flexibility of the protein and more importantly the appearance of a first partial opening of the pincers region.

**Figure 6.**
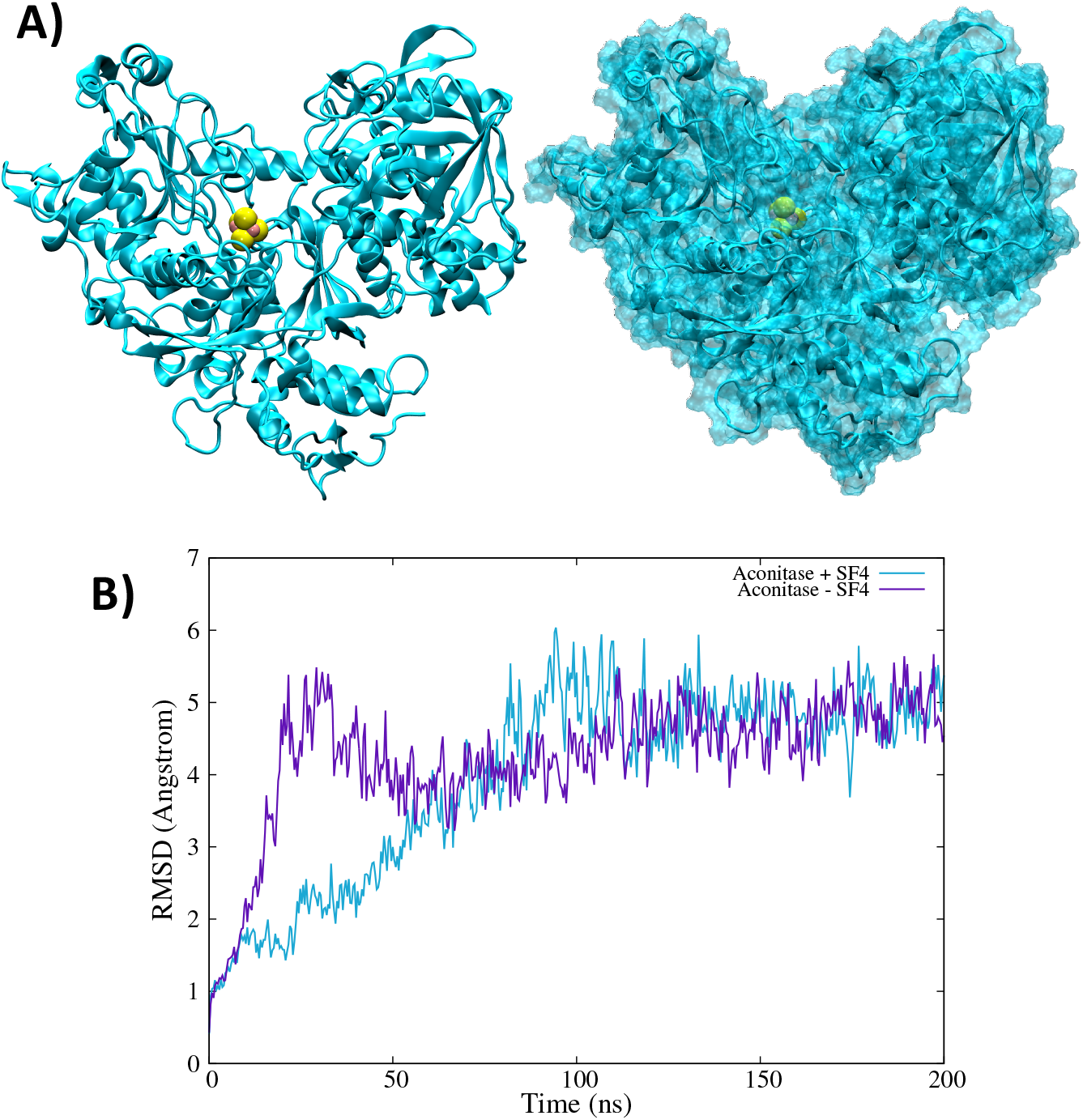
A) representative snapshots of the cAco dynamics in cartoon and surface representation. The Fe-S cluster is highlighted in van der Walls. B) Time evolution of the RMSD of the cAco MD simulation in presence and in absence of the Fe-S cluster.

### 3.4. Thermodynamics of the IRP/Aco transition

To better assess the thermodynamic and the structural mechanisms involved in the cAco/IRP1 conformational transition we resorted to enhanced sampling techniques providing the free energy profile for the interconversion between closed and open conformations in presence or in absence of the Fe-S cluster ligand. To this aim and due to the rather complexity of the transition we have used the RMSD difference between the reference conformations as a collective variable. Indeed, even if RMSD can be degenerate and hence could lead to the exploration of unphysical states, it allows to capture global deformations like those that are relevant here.

As seen from Figure 7A in the case of the apo system, i.e. the Fe-S-missing equivalent of IRP1, we can identify a rather large and flat free energy area corresponding to the absolute minimum extending roughly to the −2 to +2 Å ΔRMSD region. As also seen from the representative snapshots reported in the inlay this area corresponds to a closed conformation of the protein, which is equivalent to a IRP1 functional conformation. This result is also coherent with the observations of the equilibrium MD which point to an increase flexibility of the IRE-unbound IRP1 and a slight deviation from the reference crystal structure which is obtained from the IRP1/IRE complex. On the other hand, closed conformations can be observed at ΔRMSD of about −8 Å, which are fully consistent with the crystallographic structure of the functional cAco. However, the closed conformation for the apo system clearly does not correspond to a free energy minimum, and only a flat surface can be observed and is, hence, unstable. Furthermore, a free energy penalty of about 10 kcal/mol is associated to this conformation as compared to the open conformation.

**Figure 7.**
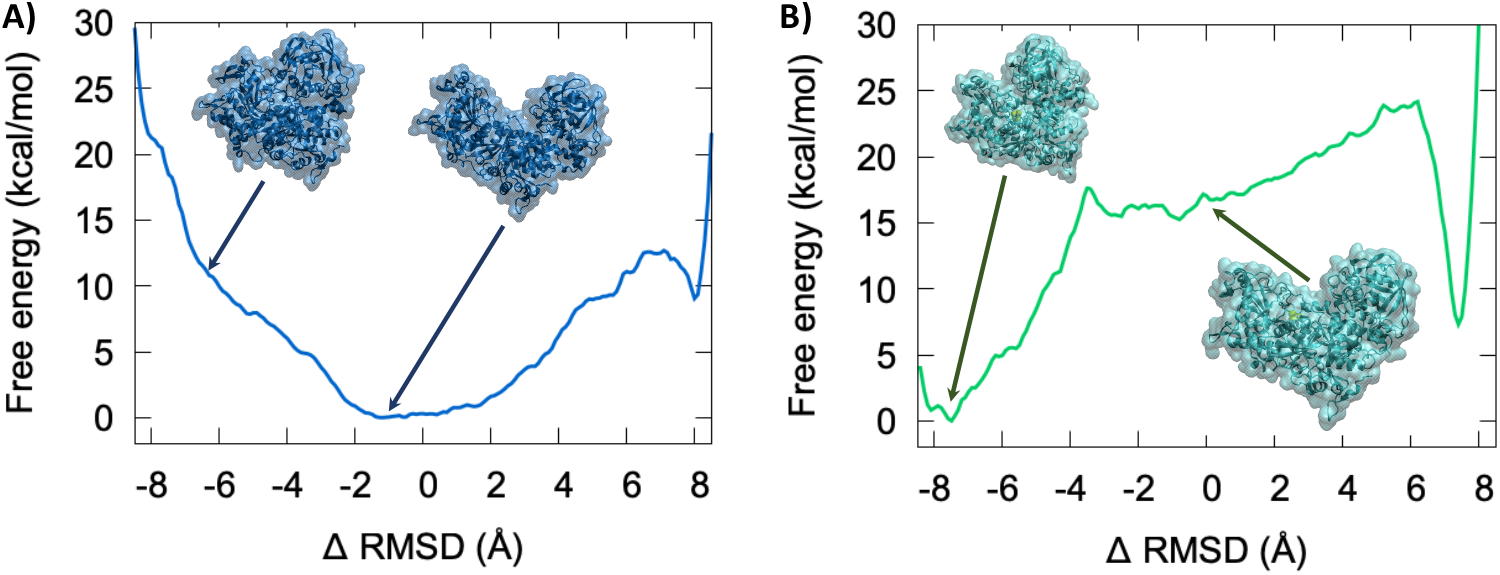
Free energy profile corresponding to the closed open (cAco/IRP1 like) transition for the apo (A) and the Fe-S cluster-containing (B) systems.

The free energy profile is drastically changed when considering the inclusion of the Fe-S cluster (Figure 7B). Indeed, while we may still recognize the flat area close to a ΔRMSD of 0.0 Å and indicative of open conformations, this area is no more associated with the global free energy minimum. The closed conformation (ΔRMSD ∼ −8.0 Å) is now highly stabilized and present a rather steep free energy well leading to a stabilization of around 15 kcal/mol compared to the open conformation. Interestingly, only a rather small barrier of about 5 kcal/mol should be overcome for passing from the open to the closed state, while the opposite transition would require bypassing 20 kcal/mol. Once again, these results are coherent with the observation raised from the equilibrium MD concerning the flexibility and the breathing mode of the apo system. The large stabilization of the closed conformation by the Fe-S cluster is mainly due to specific interactions taking place with amino acids belonging to the different protein regions which help maintaining the compact structure.

## 4. Discussion and Conclusion

By using extended MD simulations, including enhanced sampling and free energy methods, we have provided a unified vision of the molecular basis involved in the complex regulation of iron hemostasis and in the regulation of the translation of ferritin. Indeed, from the one side we have examined the specific interactions exerted between IRP1 protein and the IRE region of ferritin mRNA. Differently from other nucleic acid interacting proteins [42,58–63] IRP is able to develop strong and specific hydrogen bonds not only with the backbone but also with specific IRE nucleobases, in particular the role of the mRNA hairpin region in favoring the interaction is clearly evidenced. Since the interaction with IRP1 involves different IRE structural domains, the protein should maintain an open conformation to allow for the efficient sequestration of the mRNA. Interestingly, we have evidenced that the interaction network between IRE and IRP1 is quite complex and involves also transient interactions which are not always captured by the crystallographic structure.

The role of Fe-S cluster in driving the conformational transitions between the open (IRP1) form of the protein and its closed state (equivalent to cAco) has been rationalized. Indeed, the specific interactions developed by the Fe-S cluster, which is buried inside the protein favors the closed conformation, leading to a free energy gain of about 15 kcal/mol, only relatively small barriers (about 5 kcal/mol) should be overcome for achieving the open/closed transition. On the contrary, the open conformation is more stable in the absence of the stabilization network induced by Fe-S leading to a 10 kcal/mol preference for the open conformation. Note also that in the IRE/unbound state, the open conformation is characterized by a large and rather flat free energy surface, which is compatible with the observed breathing oscillation of the pincers.

Our results provide for the first time a clear atomistic view of the complex mechanisms at the base of FIP regulation, they are also instrumental to identify crucial regions of the IRP1/cAco protein which could be potentially druggable in the perspective of controlling iron hemostasis. The involvement of mRNA sequence in the process, and the precise determination of its structure and role, can also be potentially important in the framework of therapeutic RNA strategies, which are gaining significant momentum also in cancer therapy.

## Supplementary Materials

The following are available online at www.mdpi.com/xxx/s1, Figure S1: Time series of the RMSD for the isolated IRP1; Figure S2: Time series of the Solvent Accessible Solvent Area (SASA) for isolated IRP1 and the IRE/IRP1 complex; Figure S3: Time series of the Solvent Accessible Solvent Area (SASA) for cAco with and without the inclusion of the Fe-S cluster; Figure S4: Time series of the distances between the interfacial residues of IRP1 and IRE; Figure S5: Time series of the distance between the center of mass of the two dangling uracil nucleobases (A) and representative snapshots extracted from the MD simulation (B). Video S1: Interconversion between closed and open form of cAco in presence of the Fe-S cluster; Video S2: Interconversion between closed and open form of cAco in absence of the Fe-S cluster.

## Author Contributions

Conceptualization, A.M and S.G..; methodology, E.B., C.H.; data analysis, C.H. E.B.; writing—original draft preparation, A.M., G.H., N.T. S.G.; supervision, A.M., S.G., N.T. All authors have read and agreed to the published version of the manuscript.

## Acknowledgments

Support from the Université de Lorraine and CNRS is gratefully acknowledged. Calculations have been performed on the local LPCT computing cluster and on the Lorraine regional computing infrastructure “Explor” under the project “Dancing under the light”. E.B. thanks the CNRS ant the French Ministry of higher Education and Research (MESRI) for funding her postdoctoral position under the “GAVO” project.

## Conflicts of Interest

The authors declare no conflict of interest.

## Notes

### Competing Interest Statement

The authors have declared no competing interest.

## References

1. Abbate, V., Hider, R. Iron in biology. Metallomics 2017, 9, 1467–1469.

2. Lindley, P.F. Iron in biology: A structural viewpoint. Reports Prog. Phys. 1996, 59, 867–933.

3. Andrews, N.C. Forging a field: The golden age of iron biology. Blood 2008, 112, 219–230.

4. Beard, J.L. Iron biology in immune function, muscle metabolism and neuronal functioning. J. Nutr. 2001, 131, 568S–580S.

5. Roat-Malone, R.M. Iron-Containing Proteins and Enzymes. In Bioinorganic Chemistry; John Wiley & Sons, Ltd, 2007; pp. 343–476.

6. Fuss, J.O., Tsai, C.L., Ishida, J.P., Tainer, J.A. Emerging critical roles of Fe-S clusters in DNA replication and repair. Biochim. Biophys. Acta - Mol. Cell Res. 2015, 1853, 1253–1271.

7. Walden, W.E., Selezneva, A., Volz, K. Accommodating variety in iron-responsive elements: Crystal structure of transferrin receptor 1 B IRE bound to iron regulatory protein 1. FEBS Lett. 2012, 586, 32–35.

8. Chanet, R., Baïlle, D., Golinelli-Cohen, M.-P., Riquier, S., Guittet, O., Lepoivre, M., Huang, M.-E., Vernis, L. Fe-S coordination defects in the replicative DNA polymerase delta cause deleterious DNA replication in vivo and subsequent DNA damage in the yeast Saccharomyces cerevisiae. G3 Genes|Genomes|Genetics 2021, 11.

9. Kruszewski, M. Labile iron pool: The main determinant of cellular response to oxidative stress. Mutat. Res. - Fundam. Mol. Mech. Mutagen. 2003, 531, 81–92.

10. Chutvanichkul, B., Vattanaviboon, P., Masoodi, S., U-pratya, Y., Wanachiwanawin, W. Labile iron pool as a parameter to monitor iron overload and oxidative stress status in β-thalassemic erythrocytes. Cytom. Part B - Clin. Cytom. 2018, 94, 631–636.

11. 3 Kakhlon, O., Cabantchik, Z.I. The labile iron pool: Characterization, measurement, and participation in cellular processes. Free Radic. Biol. Med. 2002, 33, 1037–1046.

12. Cabantchik, Z.I. Labile iron in cells and body fluids: Physiology, pathology, and pharmacology. Front. Pharmacol. 2014, 5, 45.

13. Bauckman, K., Haller, E., Taran, N., Rockfield, S., Ruiz-Rivera, A., Nanjundan, M. Iron alters cell survival in a mitochondria-dependent pathway in ovarian cancer cells. Biochem. J. 2015, 466, 401–413.

14. Chen, Y., Fan, Z., Yang, Y., Gu, C. Iron metabolism and its contribution to cancer (Review). Int. J. Oncol. 2019, 54, 1143–1154.

15. Brown, R.A.M., Richardson, K.L., Kabir, T.D., Trinder, D., Ganss, R., Leedman, P.J. Altered Iron Metabolism and Impact in Cancer Biology, Metastasis, and Immunology. Front. Oncol. 2020, 10, 476.

16. Zhang, C., Zhang, F. Iron homeostasis and tumorigenesis: molecular mechanisms and therapeutic opportunities. Protein Cell 2015, 6, 88–100.

17. Porporato, P.E., Filigheddu, N., Pedro, J.M.B.S., Kroemer, G., Galluzzi, L. Mitochondrial metabolism and cancer. Cell Res.2018, 28, 265–280.

18. Gao, J., Zhou, Q., Wu, D., Chen, L. Mitochondrial iron metabolism and its role in diseases. Clin. Chim. Acta 2021, 513, 6–12.

19. Thomas, C., Mackey, M.M., Diaz, A.A., Cox, D.P. Hydroxyl radical is produced via the Fenton reaction in submitochondrial particles under oxidative stress: Implications for diseases associated with iron accumulation. Redox Rep. 2009, 14, 102–108.

20. Eisenstein, R.S., Blemings, K.P. Iron regulatory proteins, iron responsive elements and iron homeostasis. J. Nutr. 1998, 128, 2295–2298.

21. Peyssonnaux, C., Zinkernagel, A.S., Schuepbach, R.A., Rankin, E., Vaulont, S., Haase, V.H., Nizet, V., Johnson, R.S. Regulation of iron homeostasis by the hypoxia-inducible transcription factors (HIFs). J. Clin. Invest. 2007, 117, 1926–1932.

22. Winter, W.E., Bazydlo, L.A.L., Harris, N.S. The molecular biology of human iron metabolism. Lab Med. 2014, 45, 92–102.

23. Wallace, D.F. The Regulation of Iron Absorption and Homeostasis. Clin. Biochem. Rev. 2016, 37, 51–62.

24. Zhou, Z.D., Tan, E.-K. Iron regulatory protein (IRP)-iron responsive element (IRE) signaling pathway in human neurodegenerative diseases. Mol. Neurodegener. 2017 121 2017, 12, 1–12.

25. Wilkinson, N., Pantopoulos, K. The IRP/IRE system in vivo: Insights from mouse models. Front. Pharmacol. 2014, 5, 176.

26. Zhang, D.L., Ghosh, M.C., Rouault, T.A. The physiological functions of iron regulatory proteins in iron homeostasis - an update. Front. Pharmacol. 2014, 5, 124.

27. Wilkinson, N., Pantopoulos, K. IRP1 regulates erythropoiesis and systemic iron homeostasis by controlling HIF2α mRNA translation. Blood 2013, 122, 1658–1669.

28. Pantopoulos, K. Iron metabolism and the IRE/IRP regulatory system: An update. Ann. N. Y. Acad. Sci. 2004, 1012, 1–13.

29. Tong, W.H., Rouault, T.A. Metabolic regulation of citrate and iron by aconitases: Role of iron-sulfur cluster biogenesis. BioMetals 2007, 20, 549–564.

30. Gourley, B.L., Parker, S.B., Jones, B.J., Zumbrennen, K.B., Leibold, E.A. Cytosolic aconitase and ferritin are regulated by iron in Caenorhabditis elegans. J. Biol. Chem. 2003, 278, 3227–3234.

31. Khan, M.A., Walden, W.E., Theil, E.C., Goss, D.J. Thermodynamic and Kinetic Analyses of Iron Response Element (IRE)-mRNA Binding to Iron Regulatory Protein, IRP1. Sci. Rep. 2017, 7, 1–11.

32. EC, T., RA, M., H, S.-G. Structure and function of IREs, the noncoding mRNA sequences regulating synthesis of ferritin, transferrin receptor and (erythroid) 5-aminolevulinate synthase. Adv. Exp. Med. Biol. 1994, 356, 111–118.

33. Hall, K.B., Williams, D.J. Dynamics of the IRE RNA hairpin loop probed by 2-aminopurine fluorescence and stochastic dynamics simulations. RNA 2004, 10, 34–47.

34. Cairo, G., Recalcati, S. Iron-regulatory proteins: Molecular biology and pathophysiological implications. Expert Rev. Mol. Med. 2007, 9, 1–13.

35. Chen, S.C., Olsthoorn, R.C.L. Relevance of the iron-responsive element (IRE) pseudotriloop structure for IRP1/2 binding and validation of IRE-like structures using the yeast three-hybrid system. Gene 2019, 710, 399–405.

36. 3 Dupuy, J., Volbeda, A., Carpentier, P., Darnault, C., Moulis, J.M., Fontecilla-Camps, J.C. Crystal structure of human iron regulatory protein 1 as cytosolic aconitase. Structure 2006, 14, 129–139.

37. Autenrieth, F., Tajkhorshid, E., Baudry, J., Luthey-Schulten, Z. Classical force field parameters for the heme prosthetic group of cytochrome c. J. Comput. Chem. 2004, 25, 1613–22.

38. Zgarbová, M., Šponer, J., Otyepka, M., Cheatham, T.E., Galindo-Murillo, R., Jurečka, P. Refinement of the Sugar-Phosphate Backbone Torsion Beta for AMBER Force Fields Improves the Description of Z-and B-DNA. J. Chem. Theory Comput. 2015, 11, 5723–5736.

39. Dans, P.D., Ivani, I., Hospital, A., Portella, G., González, C., Orozco, M. How accurate are accurate force-fields for B-DNA? Nucleic Acids Res. 2017, 45, 4217–4230.

40. Carvalho, A.T.P., Swart, M. Correction for Electronic Structure Investigation and Parametrization of Biologically Relevant Iron-Sulfur Clusters. J. Chem. Inf. Model. 2015, 55, 1508.

41. Carvalho, A.T.P., Swart, M. Electronic structure investigation and parametrization of biologically relevant iron-sulfur clusters. J. Chem. Inf. Model. 2014, 54, 613–620.

42. Sarre, A., Stelter, M., Rollo, F., De Bonis, S., Seck, A., Hognon, C., Ravanat, J.L.J.-L., Monari, A., Dehez, F., Moe, E., et al. The three Endonuclease III variants of Deinococcus radiodurans possess distinct and complementary DNA repair activities. DNA Repair (Amst). 2019, 78, 45–59.

43. Mark, P., Nilsson, L. Structure and dynamics of the TIP3P, SPC, and SPC/E water models at 298 K. J. Phys. Chem. A 2001, 105, 9954–9960.

44. Phillips, J.C., Braun, R., Wang, W., Gumbart, J., Tajkhorshid, E., Villa, E., Chipot, C., Skeel, R.D., Kalé, L., Schulten, K. Scalable molecular dynamics with NAMD. J. Comput. Chem. 2005, 26, 1781–1802.

45. Phillips, J.C., Hardy, D.J., Maia, J.D.C., Stone, J.E., Ribeiro, J. V., Bernardi, R.C., Buch, R., Fiorin, G., Hénin, J., Jiang, W., et al. Scalable molecular dynamics on CPU and GPU architectures with NAMD. J. Chem. Phys. 2020, 153, 044130.

46. Davidchack, R.L., Handel, R., Tretyakov, M. V. Langevin thermostat for rigid body dynamics. J. Chem. Phys. 2009, 130, 234101.

47. Feller, S.E., Zhang, Y., Pastor, R.W., Brooks, B.R. Constant pressure molecular dynamics simulation: The Langevin piston method. J. Chem. Phys. 1995, 103, 4613–4621.

48. Hopkins, C.W., Le Grand, S., Walker, R.C., Roitberg, A.E. Long-time-step molecular dynamics through hydrogen mass repartitioning. J. Chem. Theory Comput. 2015, 11, 1864–1874.

49. Miyamoto, S., Kollman, P.A. Settle: An analytical version of the SHAKE and RATTLE algorithm for rigid water models. J. Comput. Chem. 1992, 13, 952–962.

50. Darden, T., York, D., Pedersen, L. Particle mesh Ewald: An N⋅log(N) method for Ewald sums in large systems. J. Chem. Phys. 1993, 98, 10089–10092.

51. Zhao, T., Fu, H., Lelièvre, T., Shao, X., Chipot, C., Cai, W. The Extended Generalized Adaptive Biasing Force Algorithm for Multidimensional Free-Energy Calculations. J. Chem. Theory Comput. 2017, 13, 1566–1576.

52. Barducci, A., Bonomi, M., Parrinello, M. Metadynamics. Wiley Interdiscip. Rev. Comput. Mol. Sci. 2011, 1, 826–843.

53. Fu, H., Zhang, H., Chen, H., Shao, X., Chipot, C., Cai, W. Zooming across the Free-Energy Landscape: Shaving Barriers, and Flooding Valleys. J. Phys. Chem. Lett. 2018, 9, 4738–4745.

54. Fu, H., Shao, X., Cai, W., Chipot, C. Taming Rugged Free Energy Landscapes Using an Average Force. Acc. Chem. Res. 2019, 52, 3254–3264.

55. Humphrey, W., Dalke, A., Schulten, K. VMD: Visual molecular dynamics. J. Mol. Graph. 1996, 14, 33–38.

56. Piccinelli, P., Samuelsson, T. Evolution of the iron-responsive element. RNA 2007, 13, 952–966.

57. Sierzputowska-gracz, H., Mckenzie, R.A., Theil, E.C. The importance of a single G in the hairpin loop of the iron responsive element (IRE) in ferritin mRNA for structure: An NMR spectroscopy study. Nucleic Acids Res. 1995, 23, 146–153.

58. Hognon, C., Garaude, S., Timmins, J., Chipot, C., Dehez, F.F., Monari, A. Molecular Bases of DNA Packaging in Bacteria Revealed by All-Atom Molecular Dynamics Simulations: The Case of Histone-Like Proteins in Borrelia burgdorferi. J. Phys. Chem. Lett. 2019, 10, 7200–7207.

59. Matoušková, E., Bignon, E., Claerbout, V.E.P., Dršata, T., Gillet, N., Monari, A., Dumont, E., Lankaš, F. Impact of the Nucleosome Histone Core on the Structure and Dynamics of DNA-Containing Pyrimidine-Pyrimidone (6-4) Photoproduct. J. Chem. Theory Comput. 2020, 16, 5972–5981.

60. Hognon, C., Monari, A. Staring at the Naked Goddess: Unraveling the Structure and Reactivity of Artemis Endonuclease Interacting with a DNA Double Strand. Molecules 2021, 26, 3986.

61. Gattuso, H., Durand, E., Bignon, E., Morell, C., Georgakilas, A.G., Dumont, E., Chipot, C., Dehez, F., Monari, A. Repair Rate of Clustered Abasic DNA Lesions by Human Endonuclease: Molecular Bases of Sequence Specificity. J. Phys. Chem. Lett. 2016, 7, 3760–3765.

62. Bignon, E., Claerbout, V.E.P., Jiang, T., Morell, C., Gillet, N., Dumont, E. Nucleosomal embedding reshapes the dynamics of abasic sites. Sci. Rep. 2020, 10, 17314.

63. Bignon, E., Gillet, N., Jiang, T., Morell, C., Dumont, E. A Dynamic View of the Interaction of Histone Tails with Clustered Abasic Sites in a Nucleosome Core Particle. J. Phys. Chem. Lett. 2021, 12, 6014–6019.

