## Supplementary Information for "The iron maiden. Cytosolic aconitase/IRP1 conformational transition in the regulation of ferritin translation and iron hemostasis"

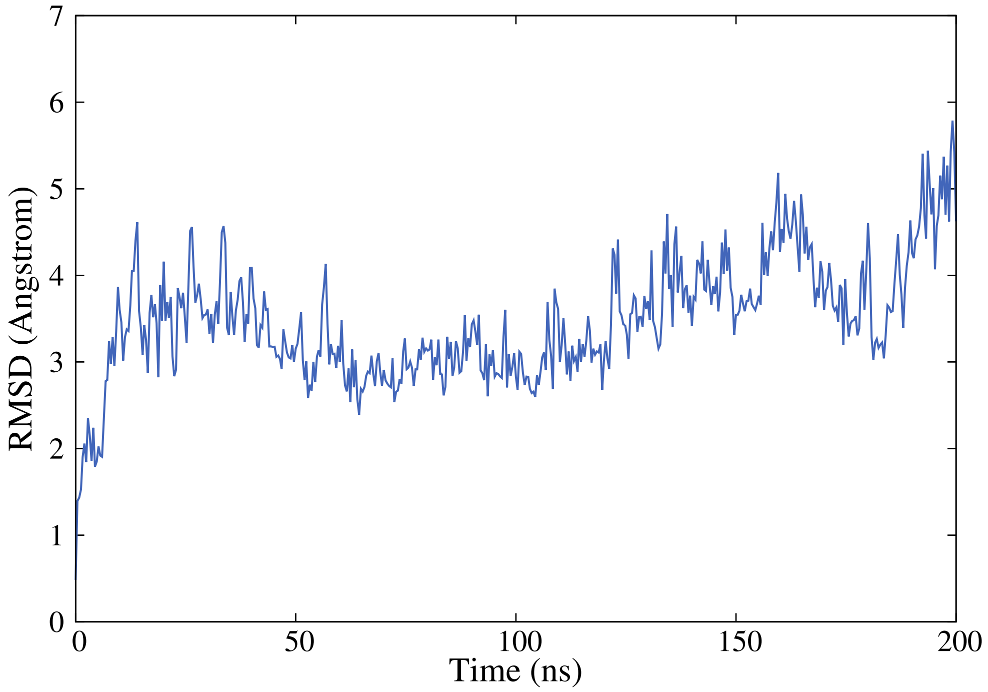


**Figure S1.** Time series of the RMSD for the isolated IRP1.

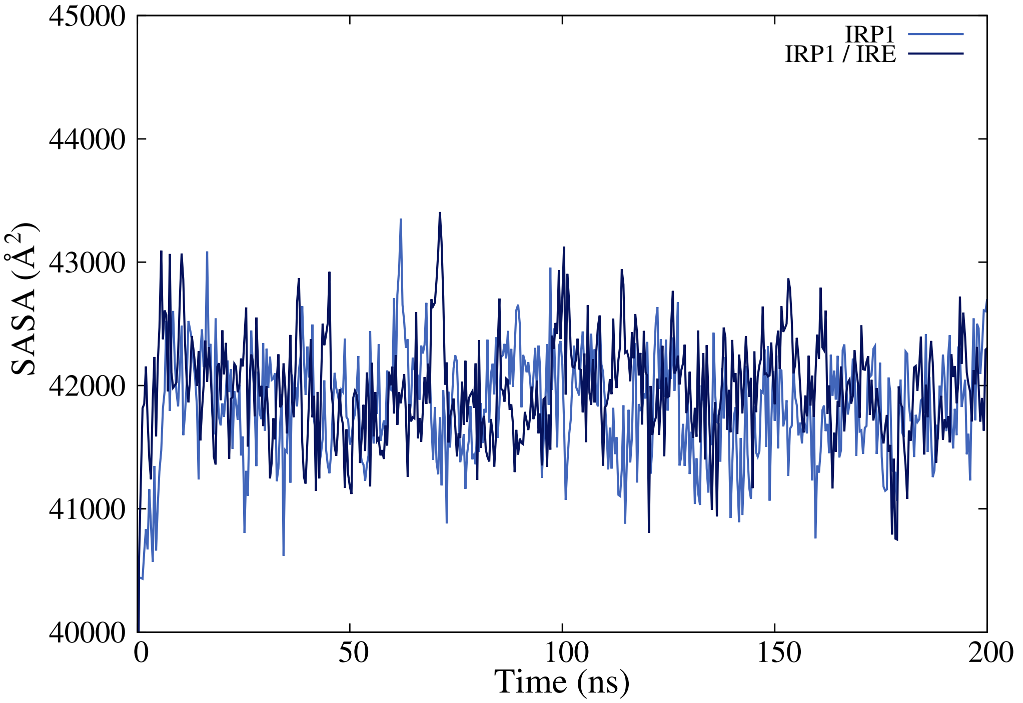


**Figure S2.** Time series of the Solvent Accessible Solvent Area (SASA) for isolated IRP1 and the IRE/IRP1 complex.

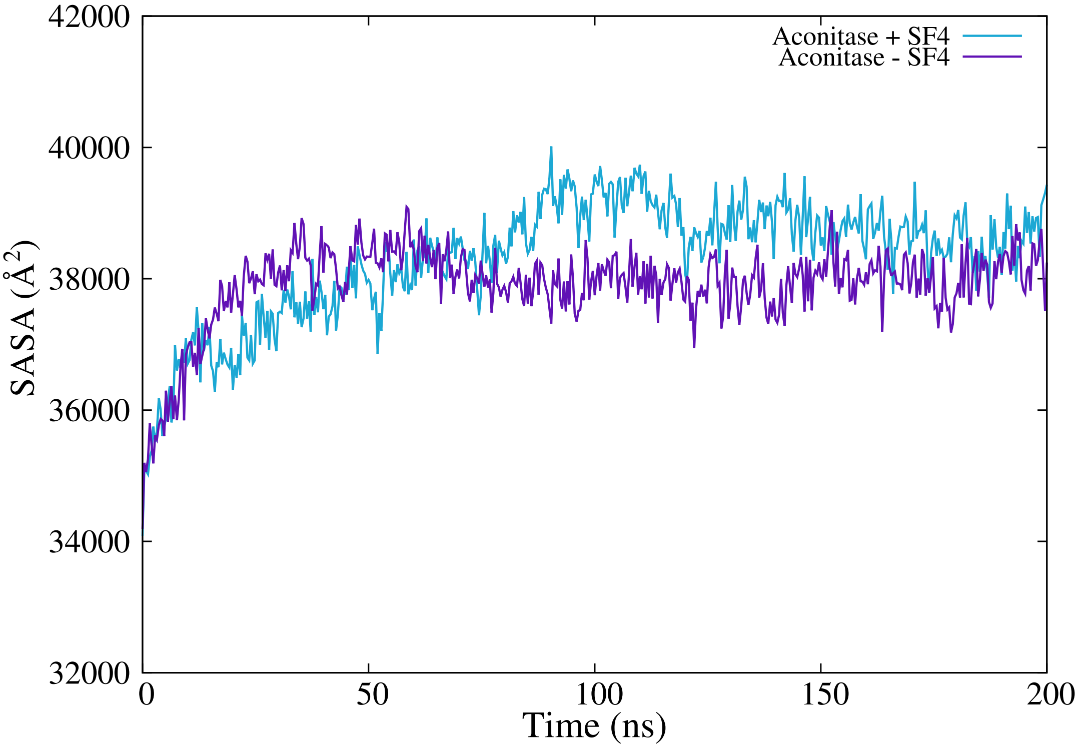


**Figure S3.** Time series of the Solvent Accessible Solvent Area (SASA) for cAco with and without the inclusion of the Fe-S cluster.

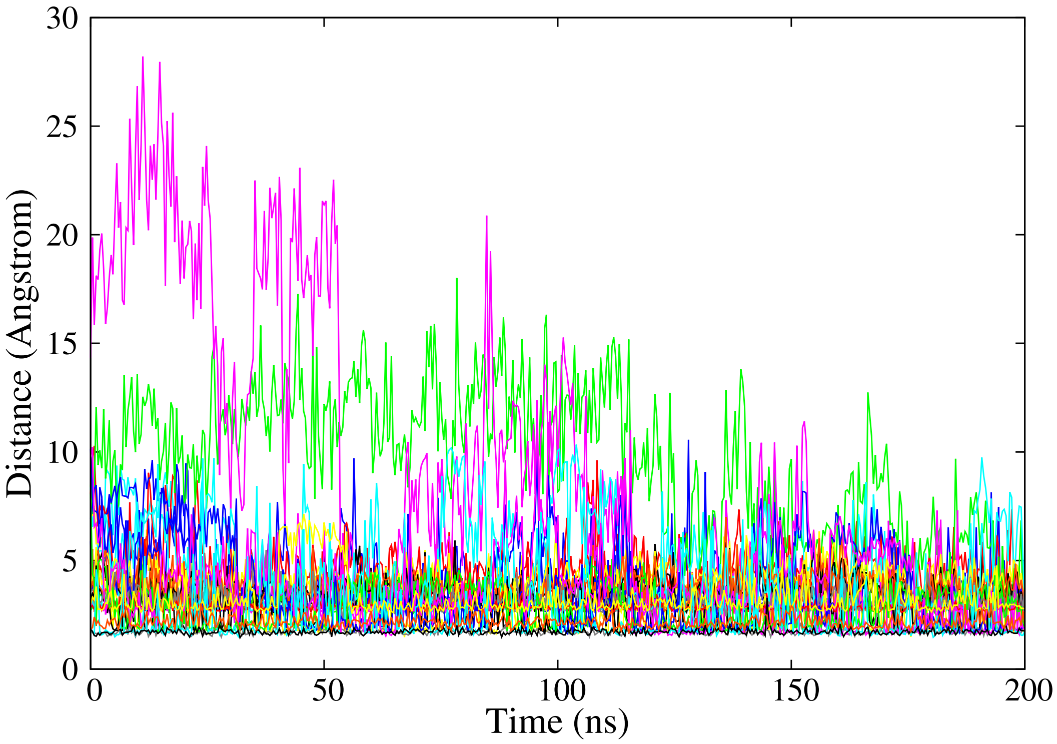


**Figure S4.** Time series of the distances between the interfacial residues of IRP1 and IRE.

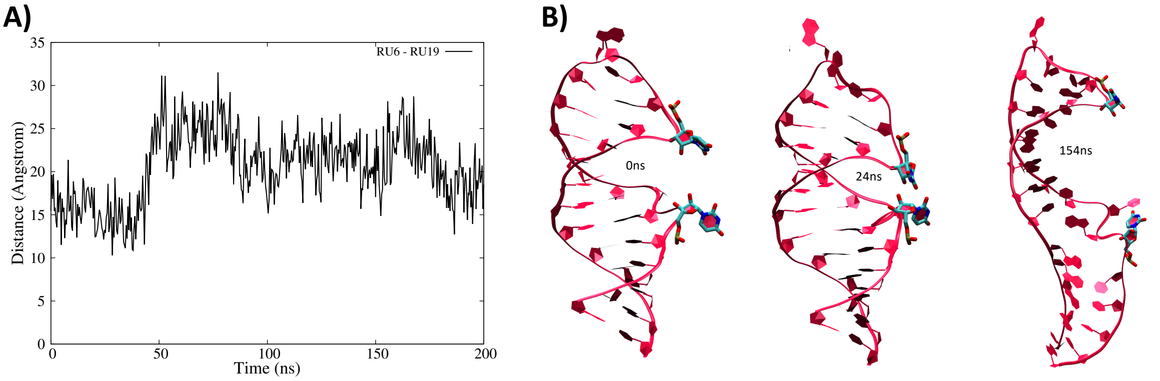


**Figure S5.** Time series of the distance between the center of mass of the two dangling uracil nucleobases (A) and representative snapshots extracted from the MD simulation (B).
